# Coverage and timeliness of vaccination and the validity of routine estimates: Insights from a Vaccine Registry in Kenya

**DOI:** 10.1101/427773

**Authors:** Ifedayo M.O. Adetifa, Boniface Karia, Alex Mutuku, Tahreni Bwanaali, Anne Makumi, Jackline Wafula, Martina Chome, Pauline Mwatsuma, Evasius Bauni, Laura L Hammitt, Christine Mataza, Collins Tabu, Tatu Kamau, Thomas N. William, J. Anthony G. Scott

**Author notes:** Corresponding author, Dr Ifedayo Adetifa, Epidemiology and Demography Department, KEMRI-Wellcome Trust Research Programme, Kilifi, PO Box 230-80108, Kilifi, Kenya, (Switchboard) (direct).

## Abstract

The benefits of childhood vaccines are critically dependent on vaccination coverage. We used a vaccine registry (as gold standard) in Kenya to quantify errors in routine coverage methods (surveys and administrative reports), to estimate the magnitude of survivor bias, contrast coverage with timeliness and use both measures to estimate population immunity.

We found coverage surveys in the 2^nd^ year of life overestimate coverage by 2%. Compared to mean coverage in infants, static coverage at 12 months was exaggerated by 7–8% for third doses of oral polio, pentavalent (Penta3) and pneumococcal conjugate vaccines, and by 24% for the measles vaccine. Surveys and administrative coverage also underestimated the proportion of the fully immunised child by 10–14%. For BCG, Penta3 and measles, timeliness was 23–44% higher in children born in a health facility but 20–37% lower in those who first attended during vaccine stock outs.

Coverage surveys in 12–23 month old children overestimate protection by ignoring timeliness, and survivor and recall biases.

## Introduction

Vaccines are the most powerful and cost effective interventions in public health; they prevent ~3 million childhood death annually, foster health equity and yield a US$44 return on investment for every US$1 spent. (1–3) However, the impact of vaccination is highly dependent on coverage. (4) Vaccine coverage estimates are widely used as a metric of performance of vaccination programmes both nationally and globally. (5–7) The benchmark of vaccination programme performance is coverage of the 3^rd^ dose of a vaccine containing diphtheria-tetanus-pertussis (DTP3) at 12 months of age. (8)

Following introduction of the Expanded Program on Immunization in 1974, global coverage of DTP3 rose to 21% by 1980. The WHO programme ‘Universal Child Immunization by 1990’ advanced this to 75% in 1990 but it remained stagnant for a further 15 years. Following the drive by WHO, UNICEF, Gavi, The Vaccine Alliance and other partners, global DTP3 coverage increased to 85% in 2015 but has stagnated again. In addition, 1-in-7 children remain unvaccinated and considerable geographical variation in coverage exists at both national and sub-national levels. (9) These factors have driven a focus on equitable access to vaccines. (6, 10)

Ideally, coverage should be measured continuously using a registry that records vaccinations received by birth cohorts, or by administrative reports.(5, 11) Vaccine registries are not routinely used especially in the low and middle income countries. So, the two principal methods supporting national and global coverage estimates are: administrative methods and random cluster surveys. Administrative methods divide the number of vaccine doses delivered by the target population estimates. Because population denominators are, on average, 5 years out of date these frequently produce estimates in excess of 100%. (12–15) Demographic and Health Survey (DHS), Multiple Indicator Cluster Survey (MICS) and EPI Cluster Surveys produce more reliable estimates because they are not dependent on census data. (16–18) However, survey methods are susceptible to selection, recall and coverage biases; for example, they fail to capture unregistered, migrant populations. (5, 19)

Coverage is typically estimated for children aged 12–23 months and referenced to their vaccination status at 12 months of age. This focuses on survivors of infancy and, if vaccination is associated with survival, there is scope for survivor bias. It also discards information on timeliness of vaccination yet it is possible that timeliness is a more sensitive indicator of health equity than a static coverage percentage. (20, 21) Finally, in a modern vaccine programme with a wide range of antigens, focus on delivery of individual vaccines does not take account of correlation between coverage of different vaccines and estimates the proportion of children who are fully immunised with difficulty.

Beyond being a performance metric, coverage is also a proxy measure of population immunity with relevance to disease control. Since coverage is closely related to disease incidence, monitoring coverage can identify likely gaps in immunity before increases in disease incidence are observed.(22) For diseases of infancy like invasive infections caused *Streptococcus pneumoniae*, coverage at 12 months of age is a poor estimate of protection during infancy and alternative measures, incorporating the timeliness of vaccination, are likely to be more useful.

Here we use a vaccine registry in Kenya (23) to quantify errors in routine coverage methods, to estimate the magnitude of survivor bias, contrast coverage with timeliness and use both measures to estimate population immunity. Finally, we examine the risk factors for delayed immunisation and illustrate the breadth of inequality in both coverage and timeliness across different birth cohorts and different locations.

## Methods

The study is an analysis of all vaccinations recorded in an electronic registry, established within the Kilifi Health and Demographic Surveillance System (KHDSS) in Kenya, between 2010 and 2016.

### Study setting and population

The KHDSS is located on the Indian Ocean coast of Kenya and comprises 280,000 residents. (24) The Kilifi Vaccine Monitoring System was established in 2009 in all 21 vaccine clinics in and around the KHDSS area. As the vaccine service expanded, we incorporated a further 15 clinics between 2011–26. (23)

As previously described, children who are residents of KHDSS, attending clinics for vaccination, are matched electronically to the KHDSS population register and all the vaccinations they received are recorded at the clinic in real time.(23, 24) Data are entered using laptop computers at the vaccine clinics and are synchronised to a bespoke MySQL v5.6.19 relational database at the main facility through weekly hard-copy transfers on laptops. Data management procedures are described elsewhere. (23, 24)

### Statistical analysis

Demographic event data (births, deaths and migrations) were combined with vaccination data from the vaccine clinics to create individual life histories of a rolling cohort of children. The data were analysed using survival analysis tools and presented as inverse survival curves with age as analysis time. We focused on two time points for estimating coverage: (A) Vaccine coverage, also referred to as ‘Up-to-date vaccination coverage’ was defined as the proportion of children vaccinated by their first birthday; (b) Age-appropriate vaccination was defined as the proportion of children vaccinated within 4 weeks of the age of vaccine eligibility in the Kenya routine childhood immunisation schedule (see table 1). (25) According to this programme, a Fully Immunized Child was defined as one who had received a dose of Bacille Calmette-Guérin vaccine (BCG), a 3-dose course of each of Pentavalent vaccine (targeting Diphtheria, Pertussis and Tetanus [DPT], Hepatitis B and *Haemophilus influenzae* b), Oral Polio Vaccine (OPV) and Pneumococcal Conjugate Vaccine (PCV), and a first dose of measles-containing vaccine (MCV1) before his or her first birthday. (26)

**Table 1.** The Kenyan childhood immunisation schedule^*^.

| **Vaccine** | **Programme started** | **Birth** | **6 wks** | **10 wks** | **14 wks** | **9 mths** | **18 mths** |
| --- | --- | --- | --- | --- | --- | --- | --- |
| BCG |  | **1** |  |  |  |  |  |
| Oral Polio Vaccine (OPV) |  | **0** | **1** | **2** | **3** |  |  |
| Inactivated Polio Vaccine (IPV) | Nov 2015 |  |  | **1** |  |  |  |
| DPT-HepB-Hib (Pentavalent) | Nov 2001 |  | **1** | **2** | **3** |  |  |
| Pneumococcal vaccine (PCV10) | Jan 2011 |  | **1** | **2** | **3** |  |  |
| Rotavirus vaccine | July 2014 |  | **1** | **2** |  |  |  |
| Measles and Rubella (MR) | June 2016 |  |  |  |  | **1** | **2** |

We sampled the event data in different ways to simulate different field approaches: (i) Birth cohort analyses. For these analyses, the denominator was the number of children who survived to the age of vaccine eligibility; the numerator was the number of these children who were then vaccinated within four weeks (age appropriate vaccination) or before their first birthday (vaccine coverage), regardless of whether they survived to these age milestones. (ii) Cross-sectional coverage surveys. We sampled the vaccine coverage status of all resident children aged 12–23 months on 1^st^ July each year. Although we used a total population sample, this still mimics the approach of a cluster sample survey. To ensure the populations in the survey analyses were linked to those in birth cohort analyses, we offset the annual birth cohorts by six months (Figure 1A); for example the 2010–11 birth cohort consisted of all children born between 1^st^ July of 2010 to 30^th^ June of 2011 and the cross-sectional survey of coverage of 12–23 month-olds, corresponding to this cohort, was undertaken on 1^st^ July 2012.

**Figure 1.**
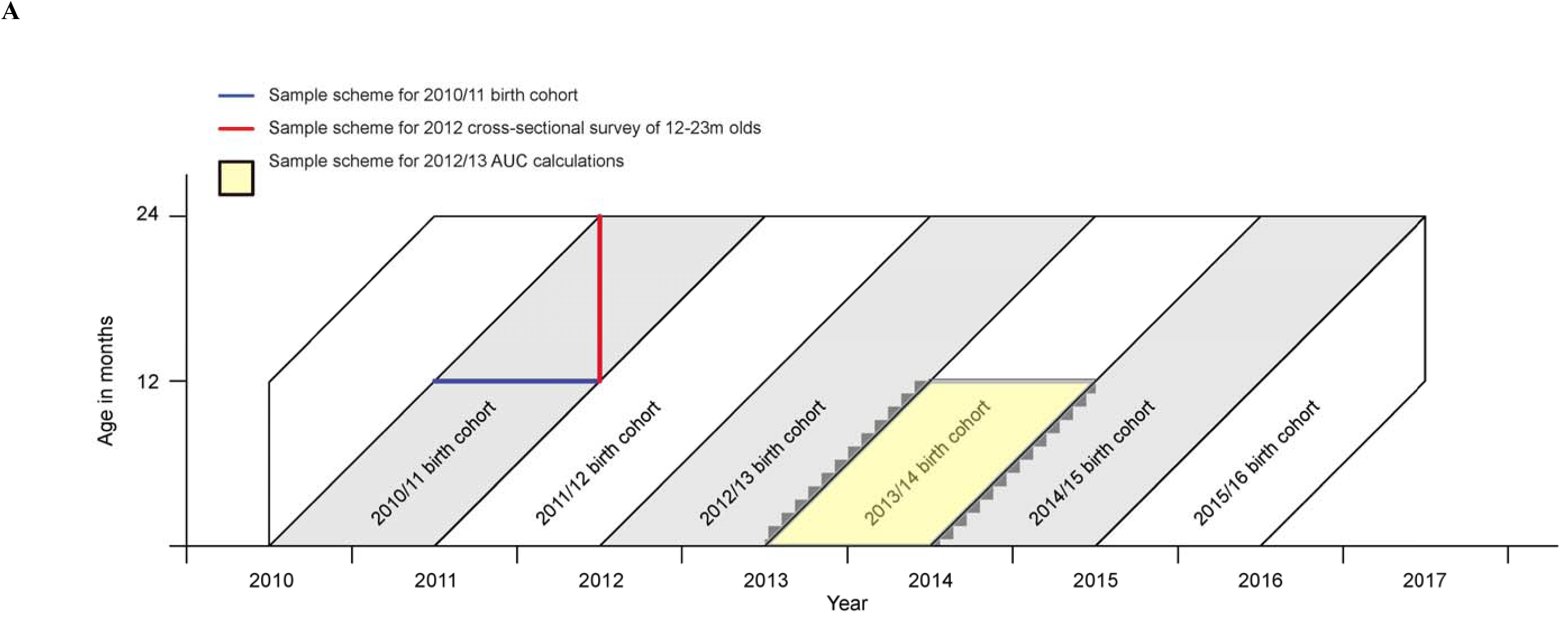
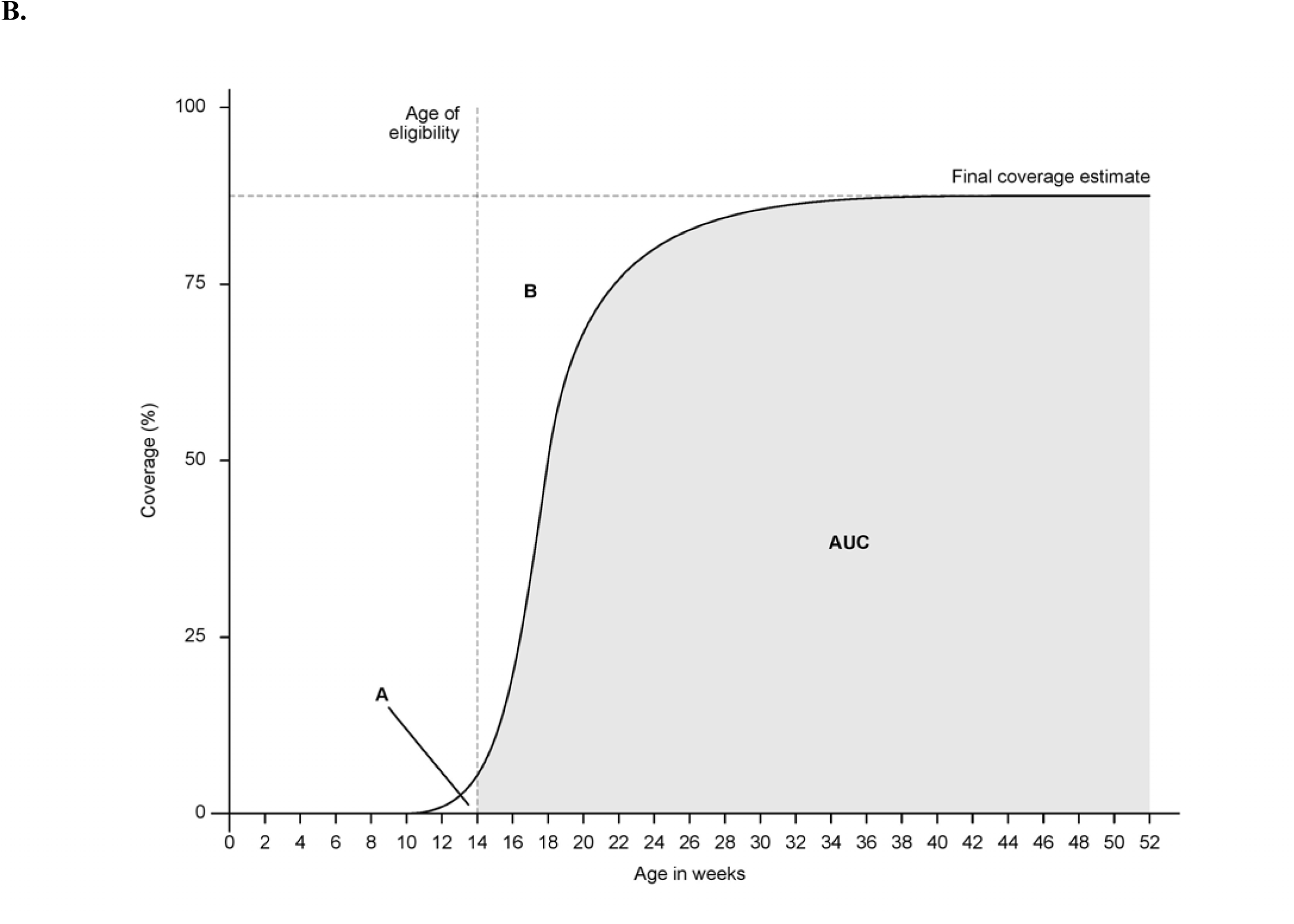
A-B- Schema showing sampling of populations for estimating vaccination coverage and parameters for area under the curve measurements

We estimated the median age at vaccination from inverse survival curves for each vaccination type. We estimated the timeliness of vaccination as the proportion of all children vaccinated by the age of 12 months who had received their vaccine within 4 weeks of becoming age-eligible. As an indicator of population immunity, we also estimated mean vaccine coverage among eligible infants as the area under the inverse survival curves (AUC) for vaccination between the age-eligible thresholds and 12 months of age (Figure 1B). (27) Age-eligible thresholds for Penta 1, 2 and 3 were 6, 10 and 14 weeks, respectively. Similar thresholds were applied to OPV and PCV. The threshold for MCV1 was 36 weeks. We also explored health equity in timeliness and final coverage over time and place by plotting inverse survival curves for BCG, Penta3 (DPT3) and MCV1 for each birth cohort and in each of the 15 administrative locations in KHDSS.

We used Cox regression models to estimate the risk factors for vaccination. The risk factors examined were drawn from variables available within the KHDSS, including place of birth, sex, birth order, maternal age and distance of the child’s home to the nearest vaccine clinic. To understand the differences between population-based and administrative coverage estimates we compared the KHDSS survey estimates for the 12–23 month old survey population in 2014 against the coverage estimates from Kilifi County in the 2014 DHS survey (28) and routinely reported administrative estimates from Kilifi County in 2014 (Kilifi County Reports). All analyses presented here are confined to children born into, and continuously resident in, the KHDSS. Coverage are presented with 95% confidence intervals.

All statistical analyses were undertaken in Stata/IC™ 13.1 (StataCorp, College Station, Texas, USA).

### Ethics statement

This study was approved by the Kenya Medical Research Institute’s (KEMRI) Scientific and Ethics Review Unit (SSC 1433).

## Results

In 6 birth cohorts (2010/11-2015/16), we studied 45 576 person years of observation (pyo) among 49 090 infants. The 6 related cross-sectional samples of children aged 12–23 months (2012–2017) comprised a total study population of 48 025. The size of each of the birth cohorts at each vaccination point in the childhood schedule, taking account of losses due to mortality and migration, is given in Figure 2.

**Figure 2.**
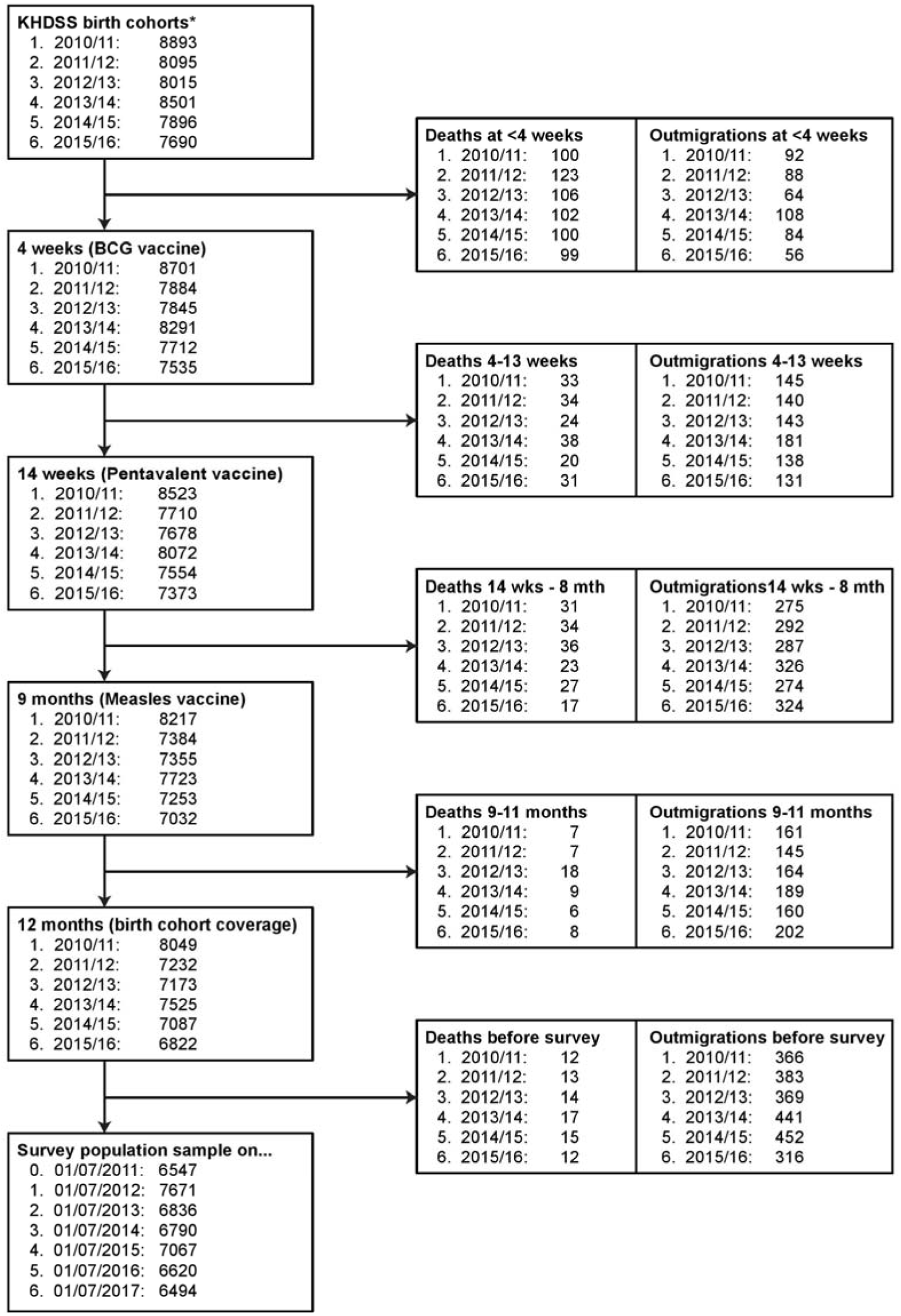
Timeline and Pentavalent vaccine (DPT3) coverage experience for annual birth cohorts compared to survey samples of 12–23-month-olds

### Vaccination coverage by birth cohort and survey

For each vaccine, coverage at age 12 months increased steadily with each advancing birth cohort but then declined in 2015–16, the last year (Table 2). The timeliness of vaccination also improved over time and then declined, though the patterns were not as consistent from vaccine to vaccine. The greatest disparity between timely coverage (4 weeks after the child became age-eligible) and vaccine coverage (at 12 months) was seen for measles vaccine, which is also the vaccine that is scheduled closest to 12 months of age. In the last 3 years of the study the estimates for coverage of the Fully Immunised Child were within 1% of the coverage estimates for measles, suggesting that the great majority of children presenting for MCV1 are already up-to-date on all other vaccines.

**Table 2.** Vaccination coverage, timeliness of vaccination and the Fully Immunised Child among residents of the Kilifi Health and Demographic Surveillance System (KHDSS) by birth cohort and survey population aged 12–23 months.

| Vaccine | Cohort | Coverage birth cohort<br>(95% CI) | Median age (weeks) | Timely<br>vaccination<br>(%)* | AUC**<br>(%) | Year | Coverage 12-23 months<br>(95% CI) | Median age (weeks) | Timely<br>vaccination<br>(%)* |
| --- | --- | --- | --- | --- | --- | --- | --- | --- | --- |
| **BCG** |  |  |  |  |  |  |  |  |  |
|  | 2010-11 | 88.7 (88.1-89.4) | 2.6 | 62.3 | 86.3 | 2012 | 91.0 (90.4-91.7) | 2.6 | 63.9 |
|  | 2011-12 | 88.5 (87.8-89.2) | 2.1 | 68.4 | 86.9 | 2013 | 91.2 (90.5-91.9) | 2.1 | 71.0 |
|  | 2012-13 | 89.8 (89.1-90.5) | 1.9 | 75.1 | 88.7 | 2014 | 93.4 (92.8-93.9) | 1.7 | 78.8 |
|  | 2013-14 | 92.6 (92.0-93.2) | 1.0 | 85.8 | 92.1 | 2015 | 96.2 (95.7-96.6) | 1.0 | 89.5 |
|  | 2014-15 | 94.2 (93.7-94.7) | 0.6 | 87.4 | 93.8 | 2016 | 96.0 (95.6-96.5) | 0.6 | 89.4 |
|  | 2015-16 | 89.0 (88.3-89.7) | 0.9 | 77.8 | 88.3 | 2017 | 90.5 (89.8-91.2) | 0.9 | 79.3 |
| **OPV3** |  |  |  |  |  |  |  |  |  |
|  | 2010-11 | 87.5 (86.8-88.2) | 15.7 | 70.5 | 81.4 | 2012 | 89.6 (88.9-90.3) | 15.5 | 72.2 |
|  | 2011-12 | 87.6 (86.8-88.3) | 15.4 | 72.3 | 82.0 | 2013 | 89.9 (89.2-90.6) | 15.4 | 74.4 |
|  | 2012-13 | 88.2 (87.5-89.0) | 15.8 | 65.2 | 80.7 | 2014 | 91.4 (90.7-92.0) | 15.7 | 68.1 |
|  | 2013-14 | 91.6 (90.9-92.2) | 15.2 | 77.3 | 86.2 | 2015 | 94.5 (93.9-95.0) | 15.2 | 80.1 |
|  | 2014-15 | 90.8 (90.1-91.4) | 16.5 | 55.8 | 78.9 | 2016 | 92.1 (91.4-92.7) | 16.4 | 56.9 |
|  | 2015-16 | 85.9 (85.1-86.7) | 18.2 | 49.1 | 74.8 | 2017 | 87.0 (86.1-87.8) | 17.9 | 50.2 |
| **Penta3** |  |  |  |  |  |  |  |  |  |
|  | 2010-11 | 85.1 (84.4-85.9) | 20.5 | 35.0 | 68.1 | 2012 | 86.8 (86.0-87.5) | 20.4 | 35.6 |
|  | 2011-12 | 87.7 (86.9-88.4) | 15.5 | 68.0 | 81.2 | 2013 | 89.8 (89.2-90.6) | 15.5 | 69.8 |
|  | 2012-13 | 88.5 (87.8-89.2) | 15.7 | 67.2 | 81.6 | 2014 | 91.7 (91.1-92.4) | 15.5 | 70.2 |
|  | 2013-14 | 91.7 (91.1-92.3) | 15.2 | 78.1 | 86.6 | 2015 | 94.6 (94.1-95.1) | 15.2 | 80.8 |
|  | 2014-15 | 93.0 (92.4-93.6) | 14.9 | 82.9 | 88.7 | 2016 | 94.6 (94.1-95.1) | 14.9 | 84.8 |
|  | 2015-16 | 87.2 (86.5-88.0) | 14.9 | 80.3 | 83.8 | 2017 | 88.6 (87.8-89.4) | 14.9 | 81.8 |
| **PCV3** |  |  |  |  |  |  |  |  |  |
|  | 2010-11 | 84.0 (83.2-84.8) | 20.9 | 41.1 | 66.0 | 2012 | 85.5 (84.7-86.3) | 20.2 | 42.3 |
|  | 2011-12 | 87.5 (86.7-88.2) | 15.4 | 74.2 | 82.4 | 2013 | 89.8 (89.1-90.5) | 15.2 | 76.2 |
|  | 2012-13 | 88.2 (87.5-88.9) | 15.7 | 67.0 | 81.2 | 2014 | 91.5 (90.8-92.1) | 15.5 | 70.0 |
|  | 2013-14 | 91.6 (91.0-92.2) | 15.2 | 77.6 | 86.4 | 2015 | 94.5 (94.0-95.1) | 15.2 | 80.4 |
|  | 2014-15 | 92.9 (92.3-93.5) | 14.9 | 82.4 | 88.6 | 2016 | 94.6 (94.0-95.1) | 14.9 | 84.4 |
|  | 2015-16 | 87.3 (86.5-88.0) | 14.9 | 79.4 | 83.7 | 2017 | 88.6 (87.8-89.4) | 14.9 | 80.9 |
| **Measles** |  |  |  |  |  |  |  |  |  |
|  | 2010-11 | 79.4 (78.5-80.3) | 41.1 | 39.6 | 57.9 | 2012 | 80.6 (79.7-81.5) | 41.0 | 39.8 |
|  | 2011-12 | 78.2 (77.3-79.2) | 41.7 | 33.5 | 53.8 | 2013 | 79.8 (78.8-80.7) | 41.6 | 34.1 |
|  | 2012-13 | 81.7 (80.8-82.5) | 41.4 | 36.4 | 57.8 | 2014 | 84.0 (83.2-84.9) | 41.0 | 37.4 |
|  | 2013-14 | 84.7 (83.9-85.5) | 40.9 | 36.4 | 59.8 | 2015 | 87.1 (86.3-87.9) | 40.7 | 37.3 |
|  | 2014-15 | 83.8 (83.0-84.7) | 41.1 | 31.0 | 57.7 | 2016 | 84.8 (83.9-85.7) | 41.1 | 31.3 |
|  | 2015-16 | 77.5 (76.6-78.5) | 41.6 | 27.8 | 52.9 | 2017 | 77.5 (76.5-78.6) | 41.6 | 27.8 |
| **FIC** |  |  |  |  |  |  |  |  |  |
|  |  |  |  |  |  | 2011 | 71.7 (70.7-72.8) | - | - |
|  | 2010-11 | 71.5 (70.6-72.5) | - | - | - | 2012 | 73.1 (72.1-74.1) | - | - |
|  | 2011-12 | 74.2 (73.3-75.2) | - | - | - | 2013 | 76.2 (75.1-77.2) | - | - |
|  | 2012-13 | 78.9 (77.9-79.8) | - | - | - | 2014 | 81.4 (80.4-82.3) | - | - |
|  | 2013-14 | 84.0 (83.2-84.8) | - | - | - | 2015 | 86.4 (85.6-87.2) | - | - |
|  | 2014-15 | 82.8 (81.9-83.6) | - | - | - | 2016 | 83.7 (82.8-84.6) | - | - |
|  | 2015-16 | 76.8 (75.8-77.8) | - | - | - | 2017 | 76.8 (75.8-77.8) | - | - |

BCG Bacille Calmette-Guérin vaccine (BCG), OPV3 Oral Polio Vaccine 3<sup>rd</sup> dose, Pentavalent Vaccine 3<sup>rd</sup> dose (Diphtheria, Pertussis, Tetanus, *Haemophilus influenzae* b and Hepatitis B combination vaccine), PCV 3 Pneumococcal Conjugate Vaccine 3<sup>rd</sup> dose, MCV1 Measles-Containing Vaccine 1<sup>st</sup> dose.
\* Proportion of vaccinated children who received their vaccines within 4 weeks of become age-eligible for vaccination
\*\* AUC % Area Under the Curve (see figure 1B)

The estimate of vaccine coverage derived from sampling children aged 12–23 months was greater than that derived from the birth cohort analysis in all birth cohorts and for all vaccines, except for measles vaccine in the 2015–16 birth cohort where the two coverage estimates were the same (77.5%, Table 2). The mean difference between survey and birth cohort coverage estimates at 12 months of age were 2.6%, 2.2%, 2.2%, 2.2%, 1.4% and 1.6% for BCG, Penta3, OPV3, PCV3, MCV1 and the fully immunised child.

The patterns of coverage of other vaccine doses (OPV1, OPV2, Penta1, Penta2, PCV1, PCV2) over time, and the disparity between coverage estimation methods were similar (Supplementary Table 1).

### Comparing published coverage estimates

Vaccination coverage estimates, referenced to 2014, differed significantly by source (Table 3). KHDSS survey coverage estimates were higher than coverage estimates for Kilifi County in the 2014 Kenya DHS report for all vaccines except BCG. Compared to the KHDSS survey estimates, administrative estimates for Kilifi County in 2014 were very similar for all vaccines except for BCG, where the administrative coverage estimate was 5% higher than the KHDSS survey estimate. Measures of the proportion of children who were fully immunised were considerably lower using both the DHS survey (71.5%) and the administrative method (67.2%) than using the survey approach in the KHDSS data (81.4%).

**Table 3.** Comparison between ‘up-to-date vaccination coverage’ estimates in Kilifi and those from other sources.

| **Vaccine** | **KHDSS survey**<br>**(12-23 months) 2014** | **Kilifi County**<br>**DHS 2014(28)** | **Kilifi County DHS –**<br>**KHDSS survey** | **Kilifi County**<br>**Administrative 2014\*** | **Kilifi County**<br>**Administrative**<br>**– KHDSS survey** |
| --- | --- | --- | --- | --- | --- |
| BCG | 93.4 (92.8-93.9) | 94.3 (89.3-97.6) | 0.9 | 98.4 (98.2-98.6) | 5.0 |
| OPV3 | 91.4 (90.7-92.0) | 87.5 (81.0-92.4) | -3.9 | 92.6 (92.2-93.0) | 1.2 |
| Penta3 | 91.7 (91.1-92.4) | 84.7 (77.8-90.2) | -7.0 | 91.4 (90.9-91.8) | -0.3 |
| PCV3 | 91.5 (90.8-92.1) | 87.4 (81.0-92.4) | -4.1 | 91.9 (91.4-92.3) | 0.4 |
| MCV1 | 84.0 (83.2-84.9) | 83.7 (76.2-89.0) | -0.3 | 84.0 (83.4-84.6) | 0 |
| FIC* | 81.4 (80.4-82.3) | 71.5 (63.4-78.7) | -9.9 | 67.2 (66.4-76.9) | -14.2 |

\*Obtained from routine immunisation reports by Kilifi County Department of Health, Kilifi, Kenya

### Timeliness of vaccination

The proportion of children vaccinated by 12 months of age who had received the vaccine within 4 weeks of becoming eligible (‘Timely vaccination’) is presented in Table 2 and Supplementary Table 1. The proportion of vaccinated infants receiving a timely vaccination increased over time for BCG, Penta3 and PCV3 but declined for OPV3 and MCV1.

Predictably, the mean coverage among eligible infants i.e. the area under the curve (AUC) was always lower than the final vaccine coverage at 12 months. The difference between these two estimates was 1.1% for BCG, 7.9% for OPV3, 7.2% for PCV3 and Penta 3 and 24.2% for MCV1. These differences are reflected in the spread of the inverse survival curves in Figure 3. Apart from 2010–11, which was the year of introduction of PCV3, delivery of PCV3 and Penta3 was timely, as was BCG. However, there is marked variation in the timeliness of OPV3 by birth cohort and timeliness of MCV1 is consistently poor across all birth cohorts.

**Figure 3 -.**
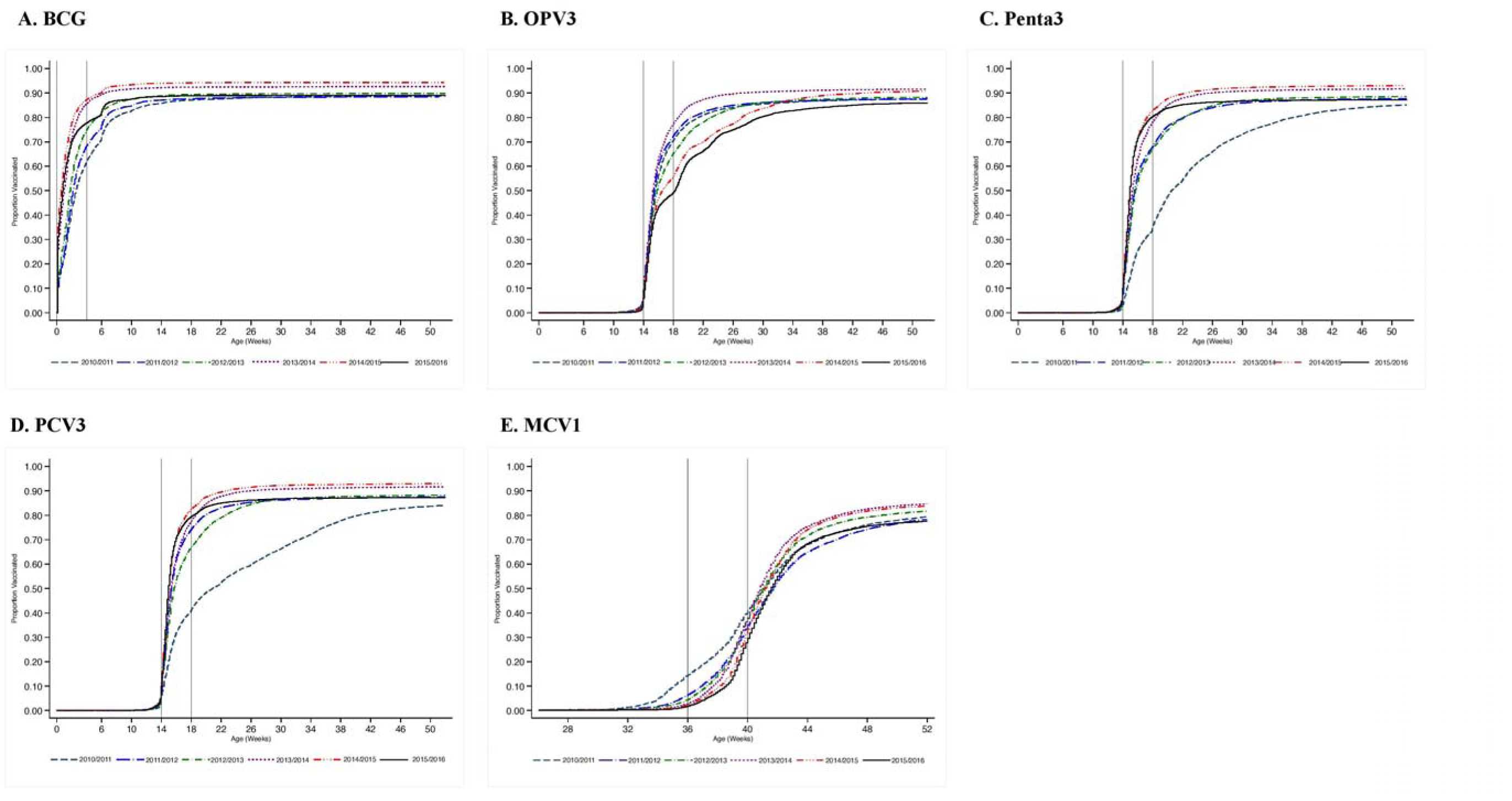
Age-specific vaccination coverage (inverse Kaplan-Meier estimates) in childhood residents of the Kilifi Health and Demographic Surveillance System by birth cohort

### Inequality in timeliness and coverage by location

Variation in the inverse survival curves for vaccination with BCG, Penta3 and MCV1 by administrative location is illustrated in Figure 4. The dispersion of the curves is greatest for MCV1 (figure 4E) and least for BCG. For BCG, the AUCs varied from 86.3% to 93.8%; for Penta3, AUCs varied from 68.1% to 88.7%; and for MCV1 AUCs varied from 52.9% to 59.8%. Some inequality in timeliness of Penta3 vaccination is apparent in figures 4C and D. The take-off in the curve for MCV1, indicating the first age at vaccination, varied from location to location, with some locations starting at 33 weeks and others not beginning until 36 weeks (figures 4E and F). In these figures, age-appropriate vaccination is estimated by the y-axis value as each curve traverses the second vertical line (4 weeks after vaccination). These ranges are 68–82% (BCG), 58–78% (Penta3) and 19–49% (MCV1). ‘Up-to-date coverage’ at age 12 months varied within the ranges 84–92% (BCG), 81–91% (Penta3) and 70–82% (MCV1)

**Figure 4 -.**
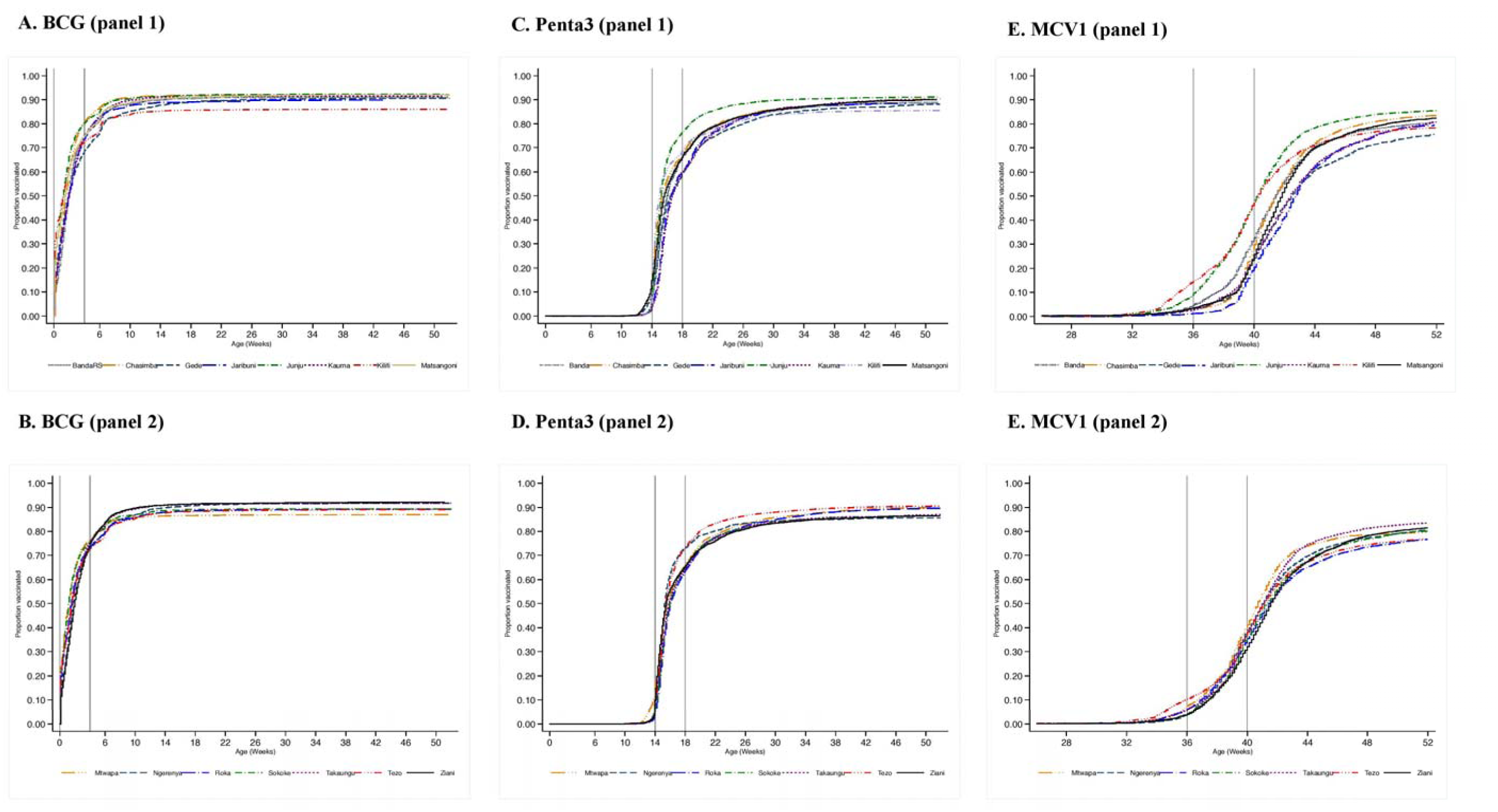
Age-specific vaccination coverage (inverse Kaplan-Meier estimates) in childhood residents of the Kilifi Health and Demographic Surveillance System by birth cohort and location

### Predictors of coverage and timeliness

The univariate and multivariable hazard ratios for risk factors for timely coverage are shown in Table 4. After accounting for the secular trend in improved coverage (BCG and Penta3), the factors associated with more timely uptake are delivery in a health facility and increasing birth order. Factors associated with delayed coverage are vaccine stock outs, increasing maternal age and increasing distance of the home from the vaccine clinic. The risk and beneficial factors were broadly similar for the three vaccines except that for measles maternal age and birth order were not associated with vaccine timeliness and the impact of stock outs was less remarkable. The results for PCV3 and OPV3 (Table S2) reflect those of Penta3.

**Table 4.** Predictors of up-to-date and age-appropriate vaccination among children in the KHDSS by birth cohort.

| Risk factors | Univariate analyses |  |  |  |  |  | Multivariable analyses** |  |  |  |  |  |
| --- | --- | --- | --- | --- | --- | --- | --- | --- | --- | --- | --- | --- |
|  | BCG |  | Penta 3 |  | MCV1 |  | BCG |  | Penta 3 |  | MCV1 |  |
|  | HR* | 95% CI | HR | 95% CI | HR | 95% CI | HR | 95% CI | HR | 95% CI | HR | 95% CI |
| Time trend (years) | 1.12 | 1.11-1.13 | 1.14 | 1.13-1.15 | 0.99 | 0.98-1.00 | 1.13 | 1.12-1.14 | 1.14 | 1.13-1.15 |  |  |
| Male sex | 0.99 | 0.97-1.01 | 1.00 | 0.98-1.02 | 0.99 | 0.98-1.01 |  |  |  |  |  |  |
| Maternal age (years) |  |  |  |  |  |  |  |  |  |  |  |  |
| <25 | - |  | - |  | - |  | - |  | - |  |  |  |
| 25-34 | 1.04 | 1.02-1.06 | 1.02 | 1.00-1.04 | 1.01 | 0.99-1.03 | 0.95 | 0.93-0.98 | 0.98 | 0.96-1.01 |  |  |
| ≥35 | 1.07 | 1.05-1.10 | 1.00 | 0.97-1.02 | 0.98 | 0.95-1.01 | 0.95 | 0.92-0.99 | 0.95 | 0.92-0.98 |  |  |
| Place of birth |  |  |  |  |  |  |  |  |  |  |  |  |
| Home | - |  | - |  | - |  | - |  | - |  | - |  |
| Health facility | 1.52 | 1.49-1.55 | 1.39 | 1.36-1.41 | 1.22 | 1.20-1.25 | 1.44 | 1.41-1.47 | 1.25 | 1.22-1.28 | 1.23 | 1.20-1.25 |
| Distance from clinic |  |  |  |  |  |  |  |  |  |  |  |  |
| <3 km | - |  | - |  | - |  | - |  | - |  | - |  |
| ≥3 km | 0.93 | 0.91-0.95 | 0.96 | 0.94-0.98 | 0.95 | 0.93-0.96 | 0.96 | 0.94-0.98 | 0.96 | 0.96-1.00 | 0.96 | 0.94-0.98 |
| Vaccine stock out | 0.60 | 0.58-0.62 | 0.56 | 0.54-0.60 | 0.78 | 0.72-0.84 | 0.63 | 0.61-0.66 | 0.72 | 0.68-0.76 | 0.80 | 0.74-0.87 |
| Birth order |  |  |  |  |  |  |  |  |  |  |  |  |
| <2 | - |  | - |  | - |  | - |  | - |  | - |  |
| 2-5 | 1.09 | 1.07-1.11 | 1.02 | 1.00-1.03 | 1.02 | 1.00-1.04 | 1.25 | 1.22-1.29 | 1.12 | 1.09-1.14 | 1.04 | 1.01-1.07 |
| >5 | 1.10 | 1.07-1.12 | 0.99 | 0.96-1.01 | 0.95 | 0.92-0.97 | 1.31 | 1.27-1.35 | 1.10 | 1.07-1.14 | 0.96 | 0.93-1.00 |

\*HR, Hazard ratios indicate the increased 'hazard' of being vaccinated among each of the risk factor categories, compared to baseline.
\*\*Adjusted for all other variables-year of birth, sex, maternal age, place of birth, distance from clinic, stockout and birth order

## Discussion

By studying an entire population sample in a Vaccine registry, we have been able to characterise the accuracy of different methods and metrics of vaccination coverage in the setting of a vaccination programme in an LMIC. The principal findings of this analysis are: (i) vaccination coverage estimates using a survey approach in the second year of life overestimate coverage by approximately 2%; (ii) compared to a total population survey in KHDSS, the cluster-survey based DHS approach in 2014 underestimated coverage of Penta3, OPV3 and PCV3 by 4–7% but the results for BCG and MCV1 were equivalent in both methods; (iii) against the same standard, the administrative methods overestimate BCG coverage by 5% but were otherwise accurate; (iv) DHS and administrative methods considerably underestimate the proportion of children who are fully immunised by 10–14%; (v) the timeliness of vaccination in this population exhibited variation both in time and location; (vi) Factors affecting coverage (‘risk factors’) were similar for each of the antigens - the largest associations with failure to vaccinate were a vaccine stock-out at the time of presentation and birth of the child outside of a hospital; (vii) Survey estimates of vaccination coverage are a poor guide to population immunity, even though they are frequently used to populate models of vaccine impact; for all vaccines except BCG, the AUC was substantially lower (7–24%) than the survey coverage.

Overall, coverage for most vaccines in Kilifi is good and has been improving throughout 2011–2016, a period of rapid introduction of new vaccines (PCV, Rotavirus, Inactivated Polio Vaccine and the combined Measles-Rubella vaccine). Samples of our registry that mimic widely utilized household surveys (MICS or DHS) tend to overestimate coverage by approximately 2%. This is inevitable if infant survival is associated with the probability of being vaccinated - either as a function of vaccine protection or as a manifestation of the ‘healthy vaccinee’ effect.(29) The infant mortality ratio is low in KHDSS (20/1000 live births in 2016); so the scope for survivor bias is much greater in settings with high infant mortality. Given the association between survival (30) and vaccination coverage, inferences based on coverage estimates from surveys of children in the second year of life should be made with more caution.

Compared against our total population survey, coverage estimates from the DHS were relatively accurate for BCG and MCV1, the first and last vaccines of the programme but under-estimated coverage of Penta, PCV and OPV by 4–7%. DHS instruments rely on evaluation of vaccine cards and if these are missing, on the recall of the parents. The first and last immunisations are probably easier to recall. In addition, the DHS sampled 144 individuals in Kilifi in 2014, whereas the vaccine registry monitored 6790 children in that year so it is possible this disparity is the consequence of a sampling error. Both sampling methods fail to identify mobile, transitory or unregistered populations: in our analysis we excluded children migrating into the area because we did not have a verifiable record of their prior vaccination history. Population movements are associated with vaccination coverage,(31) and it is likely that survey methods overestimate vaccination coverage by failing to sample transitory sub-populations. Unfortunately, we could not quantify the impact of migration on coverage estimates.

The 2014 administrative coverage estimates were remarkably close to the survey estimates, except for BCG. The positive impression this gives of administrative methods must be tempered by the fact that the ranges for administrative coverage in individual vaccine clinics in Kilifi are 20–384%, 52–144%, 52–145% and 51–144% for BCG, OPV3, Penta3 and PCV3, respectively. It may be that simple models of population growth have made accurate predictions in this area as a whole, but that parents do not always take their children to the nearest clinic; or it may be that the administrative methods were fortuitously accurate in this relatively simple 1-year comparison.

Our study shows that the prevalence of FIC is considerably underestimated by DHS-type survey methods and is grossly underestimated by administrative methods. The similarity between the coverage for MCV1 and FIC may be attributable to clinic staff following extant policy and using any vaccine visit to catch-up on missed vaccinations. Immunisation program managers and others like Gavi have advanced the FIC as a better measure of programme performance at national level than single antigen metrics like DTP3 or MCV1, and this is now the hallmark analysis for equity of access to vaccines.(26, 32) If the FIC which is better aligned with the full benefits of vaccination is to be adopted as a performance indicator, then we will need new or improved monitoring systems that are capable of linking the identity of the child across multiple records of vaccines given. Although this is theoretically possible in vaccination cards and in clinic log books, as these vaccine registry results show, it is more effectively accomplished electronically.

The inverse Kaplan Meier analyses were useful in assessing uptake and timeliness of vaccination.(27, 33) Many studies have shown marked differences in age-appropriate and ‘up-to-date coverage’ by socioeconomic status.(34, 35) In this relatively homogenous study setting, we also find significant location-specific and year-to-year variation in timeliness. Essentially, this means unless coverage is very high (>95%), it is very likely that the district or national coverage figures will conceal significant local variation. And as seen with MCV1, this becomes more relevant the later in life a vaccine is given. This could lead to substantial islands of spatiotemporal susceptibility which may ignite as, for example, unexpected measles outbreaks.

Factors associated with high coverage were younger maternal age, more previous children, and delivery in hospital. The strongest detrimental effect was vaccine stock outs, an operational challenge that immunisation programmes should be able to tackle. Place of birth and clinic visits post-delivery have been reported by us and others as important for vaccination.(36, 37) Increased contact with healthcare services is believed to increase the likelihood of vaccination via an intermediate step of health education. An alternative explanation may be differential health-seeking behaviour related to ‘healthy vaccinee’ bias. If this is true, interventions to improve vaccination coverage by increasing deliveries at health facilities may only benefit those already likely accept vaccines. Socio-economic factors also contribute to inequities in coverage and time to vaccination. (34, 35) though these are hard to characterise reliably in a relatively homogenous rural population.

In any setting with delays in vaccination, ‘up-to-date coverage’ will be a biased measure of vaccination-induced population immunity (38) and this bias, always over-estimating protection, can be as great as 24% in our survey.(38–40) AUC is a better measure because it provides an estimate of mean vaccination coverage throughout the period of risk. However, true population immunity will be lower than the AUC because a small proportion of vaccinated children will not develop an appropriate antibody response either because of operational factors (ineffective administration, inactive vaccine, heat destroyed vaccine, etc.) or host factors (immunodeficiencies) and even those who do respond adequately will not become immune until 2 weeks after a primary vaccine or 1 week after a booster. Seroepidemiological surveys provide more accurate estimates of the population fraction protected by vaccination. They can identify at-risk groups via population immunity profiles and help inform strategies to increase or sustain population immunity such as revisions of the vaccination schedule or mass campaigns.(41) Whilst the AUC may be the best epidemiological approximation of population immunity, more reliable estimates of the vaccine induced immunity can only be obtained with serological surveys.

In LMICs, vaccines account for a rapidly increasing fraction of health expenditure. Vaccine programmes are expensive but highly effective, yet their impact is critically dependent on vaccination coverage. This analysis of vaccine registry data, in a setting typical of much of sub-Saharan Africa, illustrates that methods and measures for estimating coverage are suboptimal; biased by survival to sampling date and by recall of vaccinations, incapable of revealing small area heterogeneity, unlinked and therefore unable to estimate the prevalence of the Fully Immunised Child, blind to the timeliness of vaccination and therefore to signal the gaps this produces in population immunity. Our study can be replicated across the LMICs in Africa and as part of the Comprehensive Health and Epidemiological Surveillance System (CHESS) proposed by the INDEPTH network. (42)

The methodology of measuring vaccination coverage needs to be improved on and with modern electronic record systems and serological sampling, we have the opportunity to refine our tools and make better use of the tremendous power of vaccination.

## Authors' contributions

### Ifedayo Adetifa

Contribution: Conceptualization, Formal analysis, Validation, Investigation, Methodology, Writing—original draft, Writing—review and editing

Competing interests: No competing interests declared

### Boniface Karia

Contribution: Data curation, Formal analysis, Validation, Investigation, Writing—review and editing

Competing interests No competing interests declared

### Alex Mutuku

Contribution: Data curation, Formal analysis, Validation, Writing—review and editing

Competing interests No competing interests declared

### Tahreni Bwanaali

Contribution: Supervision, Project administration, Funding acquisition, Writing—review and editing

Competing interests: No competing interests declared

### Anne Makumi

Contribution: Supervision, Project administration, Funding acquisition, Writing—review and editing

Competing interests: No competing interests declared

### Jackline Wafula

Contribution: Resources, Project administration, Writing—review and editing

Competing interests: No competing interests declared

### Martina Chome

Contribution: Validation, Investigation, Writing—review and editing

### Pauline Mwatsuma

Contribution: Resources, Project administration, Writing—review and editing

Competing interests: No competing interests declared

### Evasius Bauni

Contribution: Investigation, Methodology, Resources, Writing—review and editing

Competing interests: No competing interests declared

### Laura L Hammitt

Contribution: Funding acquisition, Investigation, Methodology, Writing—review and editing

Competing interests: No competing interests declared

### Christine Mataza

Contribution: Resources, Investigation, Supervision, Writing—review and editing

Competing interests: No competing interests declared

### Collins Tabu

Contribution: Resources, Supervision, Writing—review and editing

Competing interests: No competing interests declared

### Tatu Kamau

Contribution: Resources, Supervision, Writing—review and editing

Competing interests: No competing interests declared

### Thomas N. William

Contribution: Investigation, Methodology, Writing—review and editing

Competing interests: No competing interests declared

### J. Anthony G. Scott

Contribution: Conceptualization, Funding acquisition, Methodology, Supervision (mentorship), Writing—review and editing

## Acknowledgements

We thank the people of Kilifi County, the Minister and staff of the country Ministry of Health. This work wold not be possible without the input of the extremely hardworking field staff, data clerks, data managers, database developers and the community liaison group. This article is published with the permission of the Director of the Kenya Medical Research Institute

---

**Table S1.** Vaccination coverage and timeliness of each vaccine in residents of the Kilifi Health and Demographic Surveillance System (KHDSS) aged 12–23 months by birth cohort and survey population aged 12–23 months (other vaccine doses)

| Vaccine | Cohort | Coverage birth cohort<br>(95% CI) | Median age<br>(weeks) | Timely<br>vaccination<br>(%)* | AUC**<br>(%) | Year | Coverage 12-23 months<br>(95% CI) | Median age (weeks) | Timely<br>vaccination<br>(%)* |
| --- | --- | --- | --- | --- | --- | --- | --- | --- | --- |
| **OPV1** |  |  |  |  |  |  |  |  |  |
|  | 2010-11 | 2010-11 | 89.8 (89.1-90.4) | 6.4 | 84.4 | 87.8 | 2012 | 92.0 (91.4-92.6) | 6.4 |
|  | 2011-12 | 2011-12 | 90.0 (89.3-90.6) | 6.3 | 84.9 | 88.2 | 2013 | 92.3 (91.7-92.9) | 6.3 |
|  | 2012-13 | 2012-13 | 90.4 (89.7-91.0) | 6.4 | 82.3 | 88.1 | 2014 | 93.8 (93.2-94.4) | 6.4 |
|  | 2013-14 | 2013-14 | 92.8 (92.2-93.4) | 6.3 | 88.3 | 91.2 | 2015 | 96.0 (95.5-96.4) | 6.3 |
|  | 2014-15 | 2014-15 | 94.1 (93.6-94.6) | 6.3 | 81.6 | 91.1 | 2016 | 95.7 (95.2-96.1) | 6.3 |
|  | 2015-16 | 2015-16 | 88.5 (87.7-89.2) | 6.4 | 70.2 | 84.6 | 2017 | 89.9 (89.1-90.6) | 6.4 |
| **OPV2** |  |  |  |  |  |  |  |  |  |
|  | 2010-11 | 2010-11 | 89.2 (88.5-89.3) | 11.0 | 78.5 | 85.5 | 2012 | 91.3 (90.7-91.9) | 10.8 |
|  | 2011-12 | 2011-12 | 89.0 (88.3-89.7) | 10.8 | 79.8 | 85.7 | 2013 | 91.3 (90.6-92.0) | 10.8 |
|  | 2012-13 | 2012-13 | 89.3 (88.6-90.0) | 11.0 | 74.4 | 84.9 | 2014 | 92.7 (92.1-93.3) | 11.0 |
|  | 2013-14 | 2013-14 | 92.5(91.9-93.0) | 10.7 | 84.0 | 89.3 | 2015 | 95.5 (95.0-96.0) | 10.7 |
|  | 2014-15 | 2014-15 | 93.2 (92.6-93.8) | 11.0 | 70.7 | 87.1 | 2016 | 94.5 (93.9-95.0) | 11.0 |
|  | 2015-16 | 2015-16 | 87.6 (86.9-88.4) | 11.5 | 58.0 | 80.2 | 2017 | 88.7 (87.9-89.5) | 11.4 |
| **Penta 1** |  |  |  |  |  |  |  |  |  |
|  | 2010-11 | 2010-11 | 89.6 (88.9-90.2) | 7.0 | 67.1 | 84.2 | 2012 | 91.6 (91.0-92.3) | 6.8 |
|  | 2011-12 | 2011-12 | 89.9 (89.2-90.6) | 6.4 | 81.4 | 87.6 | 2013 | 92.2 (91.6-94.4) | 6.4 |
|  | 2012-13 | 2012-13 | 90.4 (89.8-91.1) | 6.4 | 83.2 | 88.3 | 2014 | 93.8 (93.3-94.4) | 6.3 |
|  | 2013-14 | 2013-14 | 92.9 (92.3-93.4) | 6.3 | 88.8 | 91.3 | 2015 | 96.1(95.6-96.5) | 6.3 |
|  | 2014-15 | 2014-15 | 94.2 (93.6-94.7) | 6.3 | 91.2 | 92.9 | 2016 | 95.8 (95.3-96.3) | 6.1 |
|  | 2015-16 | 2015-16 | 88.7 (87.9-89.4) | 6.1 | 86.6 | 87.6 | 2017 | 90.3 (89.5-91.0) | 6.1 |
| **Penta2** |  |  |  |  |  |  |  |  |  |
|  | 2010-11 | 88.0 (87.3-88.7) | 14.0 | 50.2 | 71.0 | 2012 | 90.0 (89.3-90.7) | 14.0 | 51.4 |
|  | 2011-12 | 89.0 (87.3-88.7) | 11.0 | 74.7 | 77.5 | 2013 | 91.3 (90.6-91.9) | 10.8 | 76.7 |
|  | 2012-13 | 89.4 (88.3-89.7) | 11.0 | 76.0 | 77.9 | 2014 | 92.9 (92.2-93.5) | 10.8 | 79.5 |
|  | 2013-14 | 92.5 (91.9-93.0) | 10.7 | 84.6 | 81.8 | 2015 | 95.5 (95.0-96.0) | 10.7 | 87.7 |
|  | 2014-15 | 93.8 (93.3-94.4) | 10.5 | 87.9 | 83.5 | 2016 | 95.3 (94.8-95.8) | 10.5 | 89.6 |
|  | 2015-16 | 88.0 (87.2-88.7) | 10.5 | 84.0 | 78.6 | 2017 | 89.5 (88.8-90.3) | 10.5 | 85.7 |
| **PCV1** |  |  |  |  |  |  |  |  |  |
|  | 2010-11 | 88.6 (87.9-89.3) | 10.1 | 49.8 | 76.2 | 2012 | 90.5 (89.8-91.1) | 9.8 | 51.3 |
|  | 2011-12 | 89.8 (89.1-90.4) | 6.3 | 85.4 | 88.1 | 2013 | 92.1 (91.5-92.7) | 6.3 | 87.7 |
|  | 2012-13 | 90.0 (89.3-90.6) | 6.4 | 82.9 | 87.8 | 2014 | 93.5 (92.9-94.1) | 6.3 | 86.5 |
|  | 2013-14 | 92.8 (92.2-93.3) | 6.3 | 88.5 | 91.2 | 2015 | 96.0 (95.5-96.4) | 6.3 | 91.9 |
|  | 2014-15 | 94.1 (93.6-94.7) | 6.3 | 91.1 | 92.9 | 2016 | 95.8 (95.3-96.3) | 6.3 | 92.9 |
|  | 2015-16 | 88.6 (87.9-89.3) | 6.1 | 86.3 | 87.6 | 2017 | 90.2 (89.5-90.9) | 6.1 | 88.0 |
| **PCV2** |  |  |  |  |  |  |  |  |  |
|  | 2010-11 | 87.1 (86.3-87.8) | 15.0 | 45.8 | 65.7 | 2012 | 88.9 (88.1-89.5) | 14.5 | 47.3 |
|  | 2011-12 | 89.0 (88.2-89.7) | 10.8 | 81.0 | 78.5 | 2013 | 91.3 (90.6-91.9) | 10.8 | 83.2 |
|  | 2012-13 | 89.1 (88.4-89.7) | 11.0 | 75.6 | 77.7 | 2014 | 92.6 (91.9-93.1) | 10.8 | 79.2 |
|  | 2013-14 | 92.4 (91.8-93.0) | 10.7 | 84.2 | 81.7 | 2015 | 95.5 (95.0-96.0) | 10.7 | 87.3 |
|  | 2014-15 | 93.8 (93.3-94.3) | 10.5 | 87.7 | 83.4 | 2016 | 95.4 (94.5-95.5) | 10.5 | 89.5 |
|  | 2015-16 | 87.9 (87.2-88.7) | 10.5 | 83.2 | 78.5 | 2017 | 89.5 (88.8-90.3) | 10.5 | 85.0 |

BCG Bacille Calmette-Guérin vaccine (BCG), OPV3 Oral Polio Vaccine 3<sup>rd</sup> dose, Pentavalent Vaccine 3<sup>rd</sup> dose (Diphtheria, Pertussis, Tetanus, *Haemophilus influenzae* b and Hepatitis B combination vaccine), PCV 3 Pneumococcal Conjugate Vaccine 3<sup>rd</sup> dose, MCV1 Measles-Containing Vaccine 1<sup>st</sup> dose.
\* Proportion of vaccinated children who received their vaccines within 4 weeks of become age-eligible for vaccination
\*\* AUC % Area Under the Curve (see figure 1B)

**Table S2.**
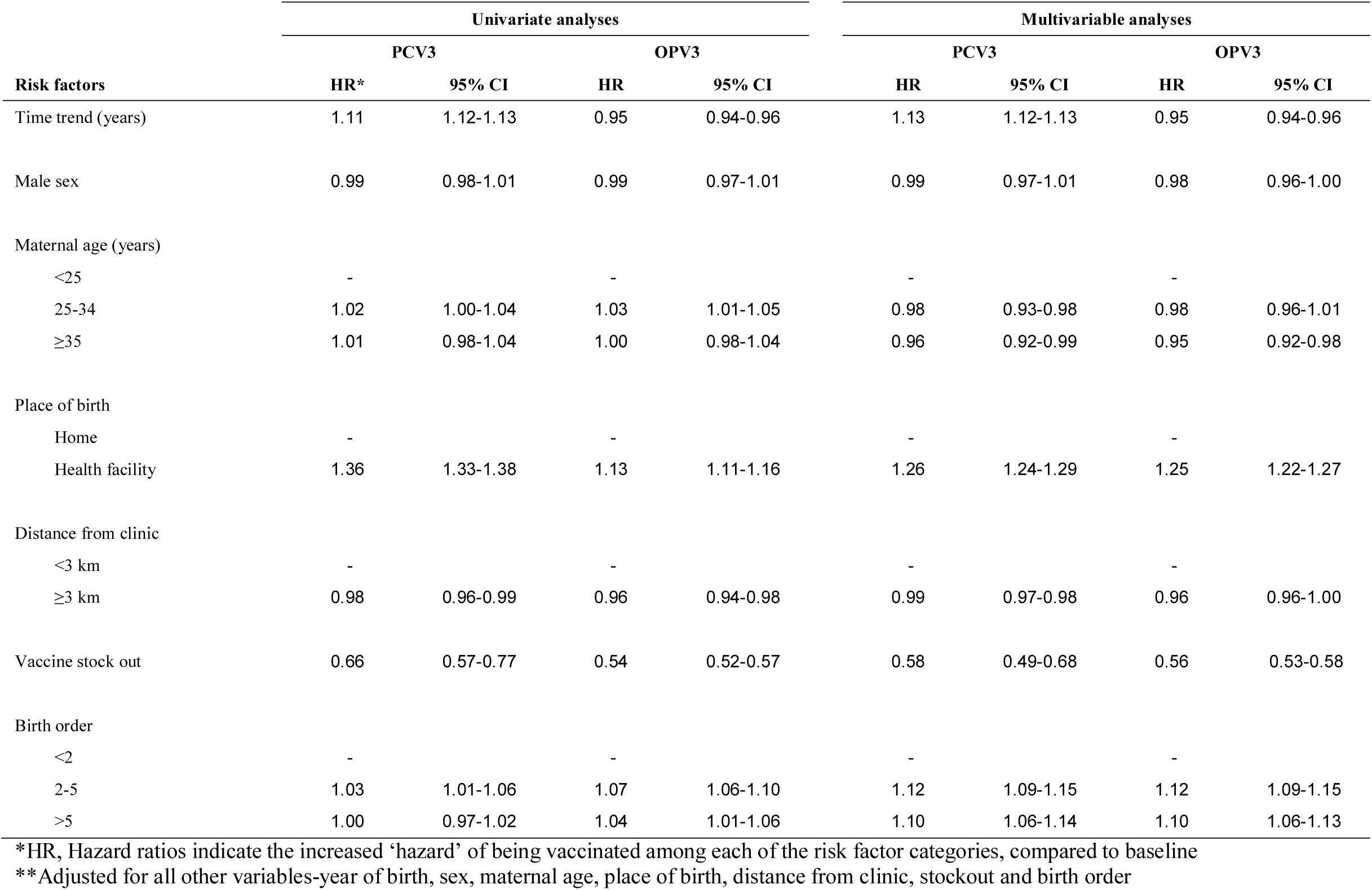
Predictors of vaccination and timeliness among children in the KHDSS by birth cohort

